# Transient oscillations of neural firing rate associated with routing of evidence in a perceptual decision

**DOI:** 10.1101/2022.02.07.478903

**Authors:** Naomi N Odean, Mehdi Sanayei, Michael N Shadlen

**Affiliations:** Zuckerman Mind Brain Behavior Institute, Department of Neuroscience, Columbia University, New York, United States; Howard Hughes Medical Institute, Columbia University, NY, USA; Kavli Institute

## Abstract

To form a perceptual decision the brain must acquire samples of evidence from the environment and incorporate them in computations that mediate choice behavior. While much is known about the neural circuits that process sensory information and those that form decisions, less is known about the mechanisms that establish the functional linkage between them. We trained monkeys to make difficult decisions about the net direction of visual motion under conditions that required trial-by-trial control of functional connectivity. In one condition, the motion appeared at different locations on different trials. In the other, two motion patches appeared, only one of which was informative. Neurons in the parietal cortex produced brief oscillations in their firing rate at the time routing was established: upon onset of the motion display when its location was unpredictable across trials, and upon onset of an attention cue that indicated in which of two locations an informative patch of dots would appear. The oscillation was absent when the stimulus location was fixed across trials. We interpret the oscillation as a manifestation of the mechanism that establishes the source and destination of flexibly routed information, but not the transmission of the information *per* se.

## Introduction

Human and animal behavior is remarkably flexible. We can execute a particular action in response to a wide variety of prompts. In a lab setting, a monkey might move its eyes to a location because a visual target had been flashed there a moment ago or because a visual stimulus at another location (or a tone) predicts a reward for this eye movement. In both scenarios there is an elevation of the firing rate of neurons that direct attention and orienting responses to the target. In the first case, the sensory input prompting this activation comes from neurons in the visual cortex with receptive fields that overlap the target location. In the second, the sensory input is from visual cortical neurons with receptive fields that do not overlap the target (or from auditory cortex). In the setting of decision making, we might say that there are many possible sources of evidence that could bear on the decision to choose a particular response. Therein lie the seeds of a routing problem that is central to cognition (***Zylberberg et al., 2010***). A general mechanism for routing is currently unknown and there is no guarantee that there is just one solution. However the study of attention and executive control implicate a role of oscillatory activity and/or synchronous spiking in the process (***Gregoriou et al., 2012**; **Lee et al., 2013**; **Saalmann et al., 2007**; **Pesaran et al., 2008**; **Dean et al., 2012**; **Stanley et al., 2018***).

We examined information routing in the context of perceptual decision-making, using a well studied direction discrimination task(***Newsome and Pare, 1988**; **Britten et al., 1992***). The subject, a rhesus monkey, must determine the net direction of dynamic random dots, only a fraction of which are informative at any moment. The decision is indicated by a saccadic eye movement to one of two choice-targets located on opposite sides of the random dot display. The decision is easy when many of the dots are moving coherently (strong motion); it is difficult when most of the dots are randomly replotted and only a small fraction of the dots are informative (weak motion). To perform well, the subject must accumulate noisy evidence over time. This accumulation is reflected in the activity of neurons in the lateral intraparietal area (LIP) with receptive fields that overlap one of the choice targets (***Shadlen and Newsome, 1996***)—an observation that presupposes a solution to a routing problem. Momentary evidence from direction selective neurons in area MT, with receptive fields that overlap the motion, must route their information, directly or indirectly, to neurons in LIP that represent the choice targets (***Salzman et al., 1992**; **Shadlen and Kandel, 2021***). This routing could not be anticipated by evolution. In some cases it might be established through learning, while in others it may need to be established on the fly. Here, we focus on the latter scenario.

We used two tasks that require a solution to the routing problem on each experimental trial. In the first, a visual cue instructs the monkey to make its decision about one of two patches of random dots *(cued attention task).* In the second, a single patch of motion appears at an unpredictable location *(variable location task).* In both tasks LIP neurons exhibit decision related activity during motion viewing, consistent with successful routing on most trials. We reasoned that the routing must be established after the onset of the attention cue or the motion stimulus and before the neurons in LIP begin to represent the accumulating evidence. We observed a prominent oscillation in the firing rates of single neurons in these epochs. The oscillation is aligned to the onset of the instructive cue in the cued attention task and to the onset of the motion stimulus itself in both tasks. The oscillations are brief and limited to the epoch preceding the representation of the accumulating evidence. We propose that they are signatures of the mechanisms that establish the routing of evidence to the site of its incorporation in a decision, but they do not appear to play a role during the information transfer accompanying decision formation.

## Results

Four rhesus monkeys *(Macaca mullata)* were trained to perform variations of the random dot motion (RDM) task that required trial by trial changes in routing. In the *cued attention task* (Fig. 1A), two patches of random dot motion were presented on each trial, preceded by a cue that indicated which location the monkey must attend to. The monkey received a reward if it chose the direction of motion in the cued patch. In the *variable location task* (Fig. 1B), just one patch of motion was presented, but its location was unpredictable.

**Figure 1.**
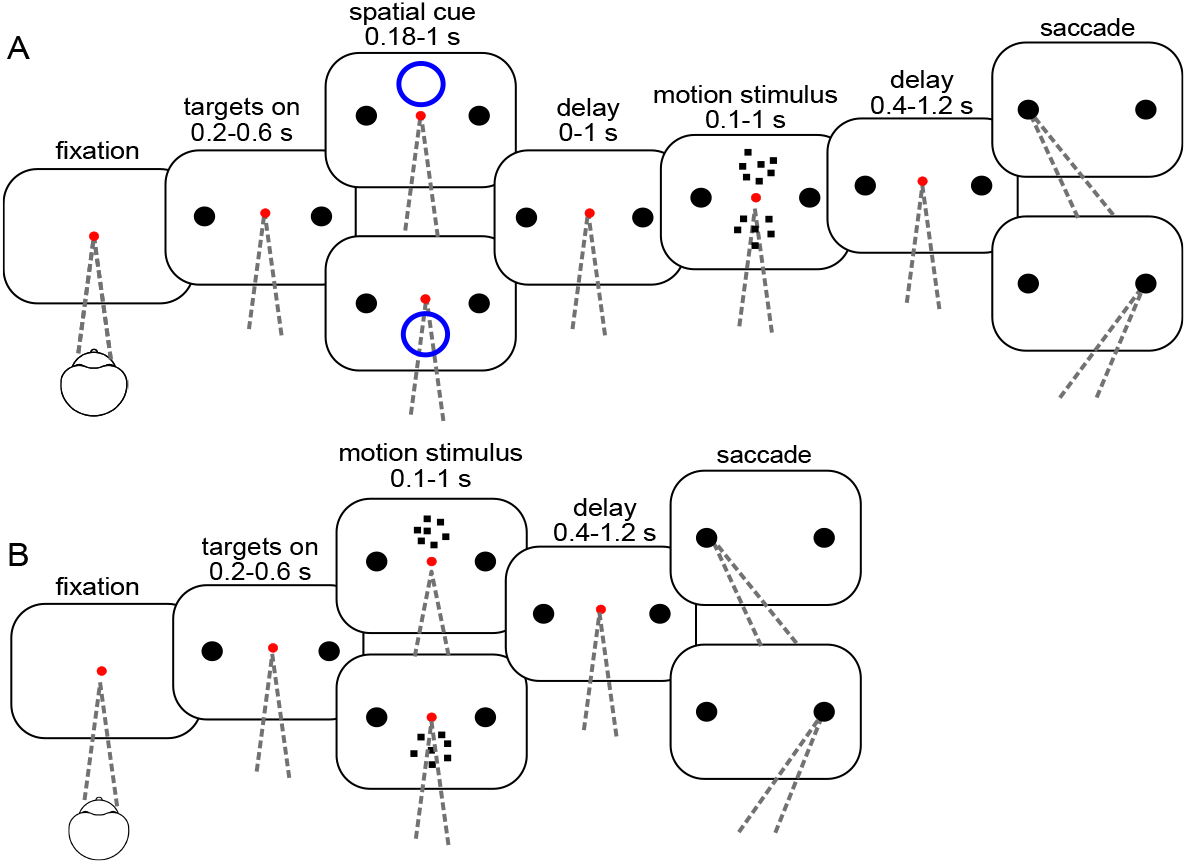
Task flow. **A**, Cued Attention task. After the monkey acquires fixation, two choice-targets appear, followed by a brief spatial cue (blue circle). After a delay, two random dot motion patches appear. The motion strength is the same in the two patches, but the directions may be the same or opposite. When the fixation point is extinguished, the monkey indicates the direction of the cued motion with a saccade to the left or right choice target and receives a reward if the choice corresponds to the direction of the cued patch. **B**, Variable Location task. Same as in *A* except there is no attention cue, and only one patch of motion is shown, either above or below the point of fixation.

In the cued attention task, both monkeys based their decisions on the relevant motion patch in at least 90% of trials (Fig. 2A) and their performance accuracy improved as a function of viewing duration at a rate consistent with temporal integration of evidence to a stopping bound, as shown previously (***Kiani et al. 2008***; Fig. 2B). However, neither monkey was able to fully ignore the uncued patch, as evidenced by the shallower choice function on the trials in which the motion patches had opposite directions (open symbols, Fig. 2A). At the strongest motion strength (± 51% coh), errors occurred on 11% of trials when the patches contained opposite directions, compared to 3% when the patches shared the same direction. Some of these errors are explained by a failure to route information from the appropriate patch. The curves in Fig. 2A&B are fits to a drift diffusion model that takes into account both motion strength and viewing duration. Importantly, it allows for the possibility that the uncued patch of dots is not fully suppressed, and on a random fraction of trials, *λ,* the monkey bases the decision on that patch (see Methods, Behavior and Table 1). In the variable location task, the monkeys made fewer than 2% errors when motion was strongest, nearly all of which were on trials with duration of less than 0.3 s (Fig. 2B). This performance is comparable to similar tasks in which the location of the motion stimulus was predictable (***Fetsch et al., 2014**; **Gold and Shadlen, 2000***). For the monkey that performed a free response version of the task, both reaction time and choice depended on the strength of motion (Fig. 2C).

**Figure 2.**
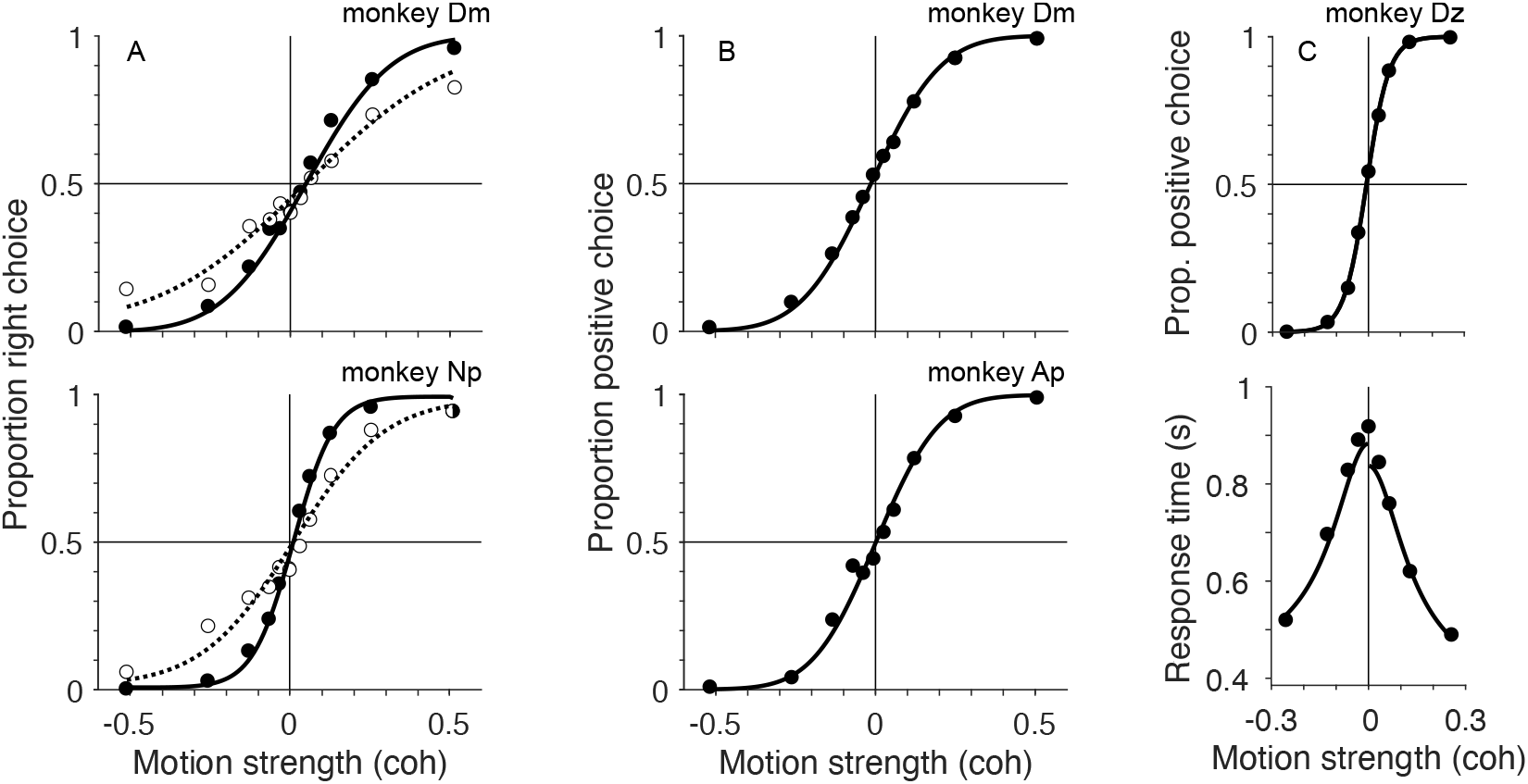
Behavior. **A**, Cued attention task. Proportion of rightward choices is plotted as a function of signed motion strength of the cued patch (positive coherence signifies rightward). Filled and open symbols show trials where the direction of uncued motion patch was the same or opposite to the cued patch, combining trials for all viewing durations. Solid and dashed curves are fits of a bounded drift-diffusion model that incorporates misrouting owing to incomplete suppression of the uncued patch or attending to it erroneously on a fraction of trials. The curves represent the expectation of the choice-proportions for the mean stimulus duration (*top*, monkey Dm; *bottom*, monkey Np). **B**, Variable location task with random stimulus durations. There is only one patch of motion. The proportion of choices in the positive direction (favoring the target in the neural response field) is plotted as as a function of signed coherence. The smooth curve is a fit to a simpler bounded drift diffusion model. As in *A*, the proportions reflect all stimulus durations, and the fit shows predictions for the mean duration (*top*, monkey Dm; *bottom,* monkey Ap). **C** Choice-response time version of the variable location task. The choices (*top*) and response times (*bottom*) are fit by a bounded drift-diffusion model. Fit parameters for all monkeys and conditions are in Table 1

**Table 1.**
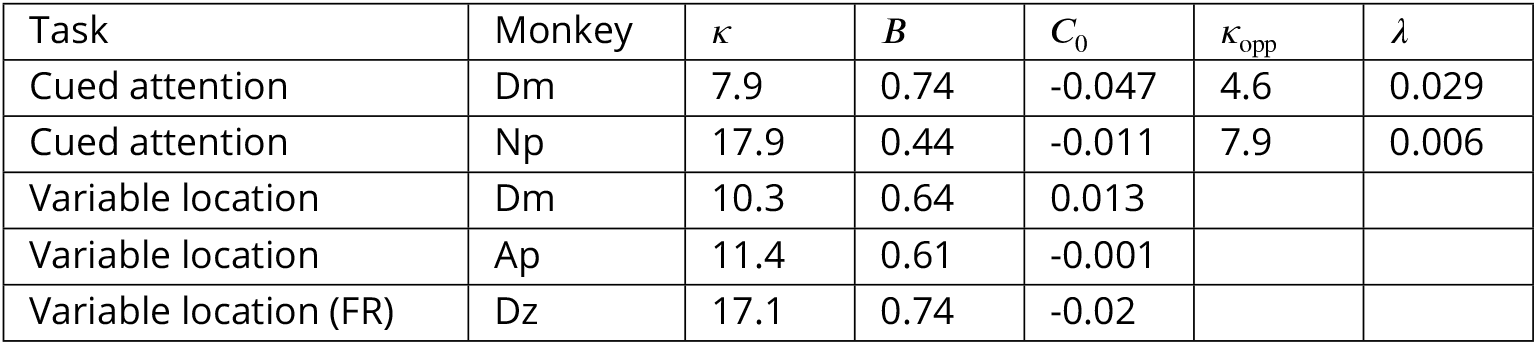
Parameters for model fits. Variables are defined in Eqs. 2, 4 and 5. Model comparison favors inclusion of ***κ***_opp_ and ***λ*** for both monkeys (BIC: 609 and 773 for Dm and NP). Inclusion of both parameters is superior to just one. For Dm, BIC = 9.7 (comparison to ***κ***_opp_ = *κ*, which attributes nearly all errors at coh = ±0.52 to use of the wrong patch). For Np, BIC = 71 (comparison to ***λ*** = 0). Free response task (FR) includes constant parameters, 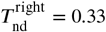 and 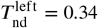, which accounts for sensory and motor latencies that add to the decision time to explain the total response time.

### Neural recordings

The data comprise 173 neurons from LIP of four monkeys (see Table 2). All neurons were screened for spatially selective persistent activity in oculomotor delayed response (ODR) tasks. One of the saccadic choice targets (*T*_in_) was placed in the neural response field, typically in the contralateral visual field (see Methods). Such neurons are known to reflect the accumulation of evidence bearing on the decision to choose *T*_in_. As shown in Fig. 3, the neural response begins to reflect the direction and strength of motion approximately 200 ms after motion onset, and this holds whether the source of evidence is from the upper or lower location. In the cued attention task, the decision-related activity is also affected by the direction of motion of the uncued patch (***Figure 3—figure Supplement 2***), consistent with the higher error rate on trials with motion in opposite directions.

**Figure 3.**
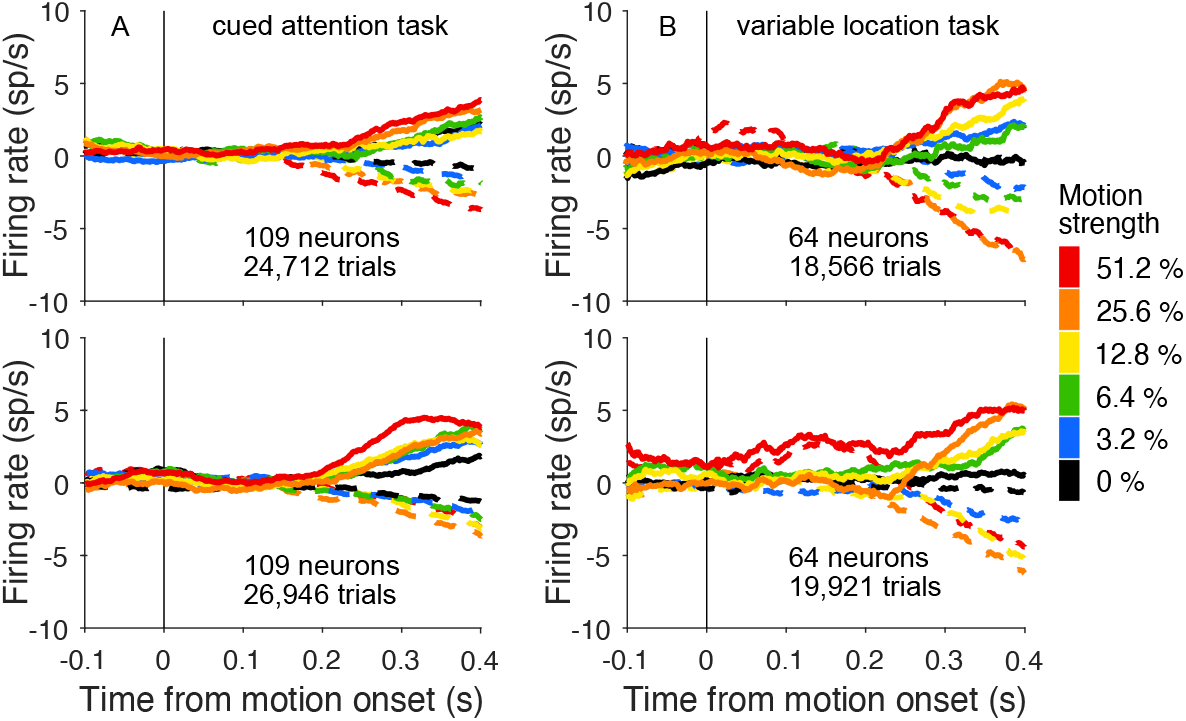
LIP neurons reflect evidence from the attended patch of motion. **A**, Activity of 109 neurons studied in the cued attention task when the upper patch *(top)* and lower patch *(bottom)* were cued as informative (combined data from monkeys Dm and Np). Responses are detrended by neuron, via subtraction of the mean firing rate, as function of time, on the lowest coherences (0% and ±3.2%). Errors on non-zero coherences are excluded. The neurons reflect the formation of decision from information derived from the upper and lower visual field. **B**, Activity of 64 neurons studied in the variable location task when the motion patch appeared in the upper *(top)* or lower *(bottom)* location (combinded data from monkeys Dm, Ap and Dz). **Figure 3—figure supplement 1.** Firing rates aligned to all task relevant events **Figure 3—figure supplement 2.** Comparison of responses when motion patches had the same or opposite directions **Figure 3—figure supplement 3.** Neural responses shown separately for each monkey and task

**Table 2.** Number of neurons recorded from each monkey in the tasks (FR, free response task; * previously published data).

|  | Np | Dm | Ap | Dz | Nt* | Br* |
| --- | --- | --- | --- | --- | --- | --- |
| Cued attention | 60 | 49 |  |  |  |  |
| Variable location |  | 28 | 23 |  |  |  |
| Variable location (FR) |  |  |  | 13 | 28 | 21 |
| Fixed location (FR) |  |  |  |  | 64 | 52 |

It thus appears that by 200 ms of the onset of motion, some mechanism must establish functional connectivity between LIP neurons that represent the choice-targets and the relevant direction-selective neurons with receptive fields that overlap the RDM. In the cued attention task, this routing might occur following the attention cue. In the variable location task, only the onset of the RDM is informative. In what follows we demonstrate a brief oscillation in the firing rates of LIP neurons. We first characterize the timing and strength of the oscillations in the two tasks. We then report additional observations that suggest the oscillations are associated with a mechanism that establishes the functional connectivity between sources of evidence and the circuits in LIP that use this evidence to establish the relative priority of the choice targets. In the Discussion, we consider how such oscillations might bear on neural mechanisms of routing.

### Oscillations in the cued attention task

Fig. 4A–D shows prominent oscillations in the firing rate of two example neurons, aligned to onset of the attention cue. Importantly, the cue, like the random dot motion, was presented outside the neural response field (see Methods, Mapping tasks). One of the neurons (Dm49) exhibits 3–4 evenly spaced periods of increased activity (~16.7 Hz) when the cue signaled that the relevant patch of motion would appear in the upper location. The other neuron (Dm35) exhibits a similar period but with a more pronounced decay in amplitude, independent of whether the cue appeared in the upper or lower location. These examples are among the most vivid in the data set. In most cases the oscillations are imperceptible in the trial rasters. The examples also highlight heterogeneous features, such as the rate of decay and spatial preference, that we will not dwell upon. What stands out as consistent is the timing, periodicity and transient nature of the oscillations. These features are preserved in the firing rate averages across the population of neurons (Fig. 4E,F).

**Figure 4.**
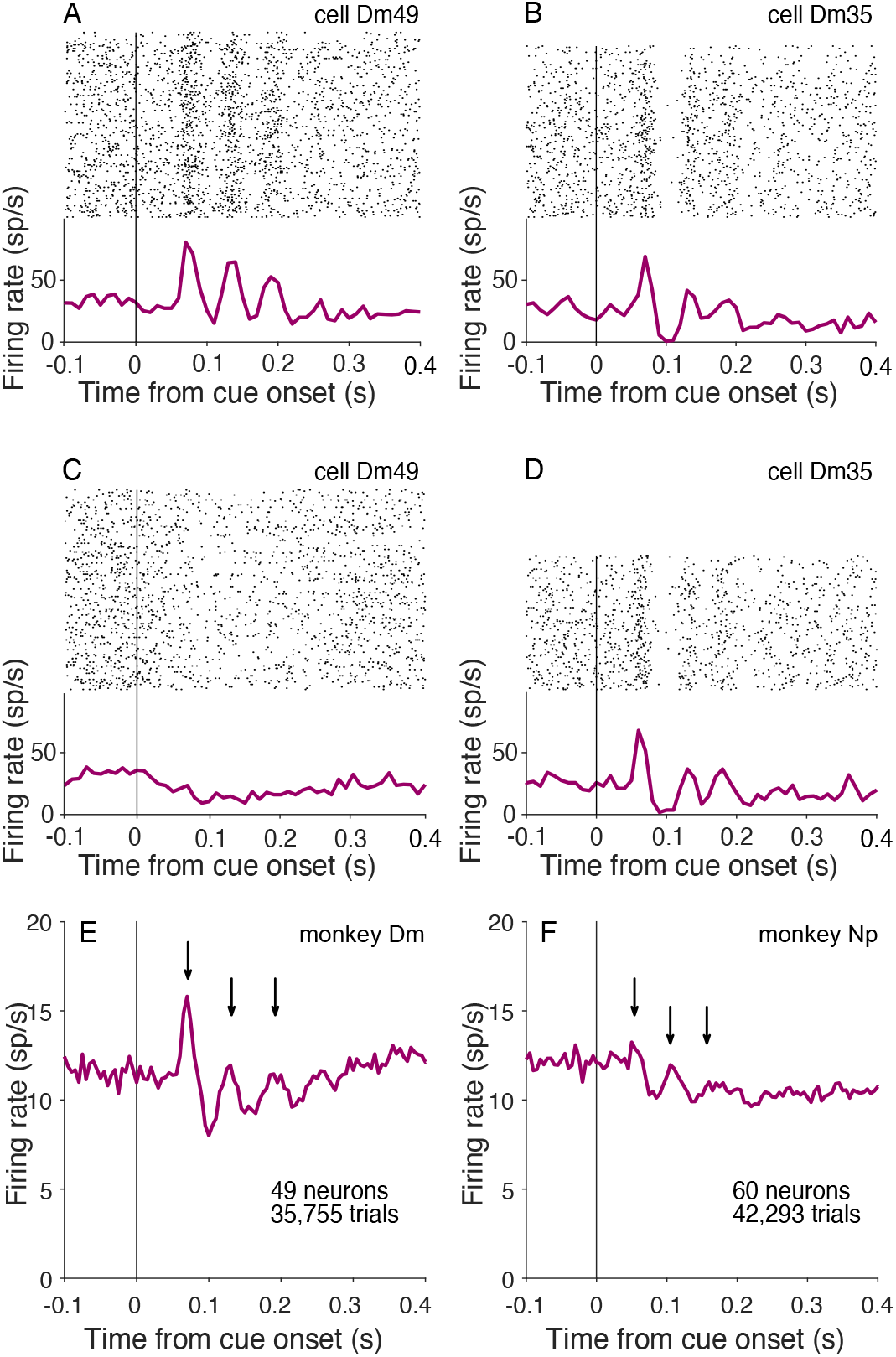
Oscillations at cue onset in the cued attention task. **A,B**, Activity of example neurons with the cue above fixation. *Top* Raster plot of spike times relative to onset of the attention cue. *Bottom* peri-event histogram shows average firing rates across trials (bin width 10 ms). **C,D**, Cue aligned activity of the same example neurons for trials with the cue below fixation. **E,F**, Average activity across all neurons combining both cue locations (bin width 5 ms). Arrows indicate peaks after motion onset. **Figure 4—figure supplement 1.** Oscillations in spiking activity measured using a matching pursuit algorithm. **Figure 4—figure supplement 2.** Realigning does not identify additional peaks

To further characterize the amplitude and frequency of these oscillations we applied a matching pursuit (MP) algorithm (***Chandran et al., 2016**; **Mallat and Zhang, 1993***) to the the across-trial average spike rate functions for each neuron. MP is especially useful for brief periodic signals, as it measures power with high temporal resolution (see Methods). We report the average Wigner-Ville power, 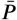, in the 12–20 Hz range using 90 ms epochs preceding and following onset of the attention cue, denoted 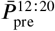 and 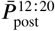 (−90 ≤ *t* < 0 and 40 ≤ *t*< 130 ms, respectively). We use the same nomenclature below, when aligning the response to other task events. The superscript identifies the range of frequencies contributing to the 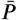 statistic. The majority of neurons recorded in the cued attention task exhibit an increase in 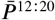 after cue onset (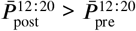, *p* < 0.05, 65 of 109 post prneurons). The presence of oscillations is similar for putative excitatory and inhibitory neurons (ascertained from spike waveform analysis; see Methods, Cell type analysis). Across the population, the mean 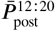 was 1.52 ± 0.49 sp^2^s^-2^, an order of magnitude larger than 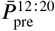 (0.12 ± 0.03 sp^2^s^-2^; *p* < 0.0001). In comparison, Wigner-Ville power in the 4–11 Hz band does not undergo change (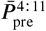 and 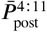 are 1.27 ± 0.29 and 1.32 ± 0.27 sp^2^s^-2^, respectively; *p* = 0.35).

The oscillation in firing rate is triggered by the onset of the cue, and decays quickly thereafter. By 0.18 s after cue onset, 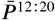 is only 0.067 ± 0.026 sp^2^s^-2^, which is comparable to 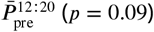 (*p* = 0.09). We wondered if the oscillations are truly brief or are simply undetectable as a consequence of dephasing. To test this, we used a piecewise linear time warp designed to realign temporally jittered oscillations *(**Williams et al., 2020**).* While this algorithm successfully realigned jittered synthetic data, it did not identify any new peaks in the neural data ***(Figure 4—figure Supplement2***). We therefore conclude that the oscillation is in fact short-lived and, in this case, caused by a task-relevant visual cue, outside the neural response field.

A weak oscillation in the firing rate is also present after motion onset. Fig. 5A–D shows the activity of the same example neurons shown in Fig. 4A–D, aligned to onset of the random dot displays. The oscillations are apparent in the rasters and average firing rates for both neurons. As shown in Fig. 5E–F, they are also evident in the average firing rate across the population of neurons. They are weaker than the oscillations induced by the attention cue (*p* < 0.0001; Fig. 5G), but they are statistically reliable: 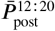 is an order of magnitude larger than 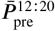 (0.80±0.30 *vs.* 0.06 ± 0.01 sp^2^s^-2^, *p* < 0.0001). The weaker oscillation following motion onset is consistent with the hypothesis that these oscillations play a role in establishing functional connectivity. In the cued attention task, information about the location of the relevant motion patch was already supplied by the cue.

**Figure 5.**
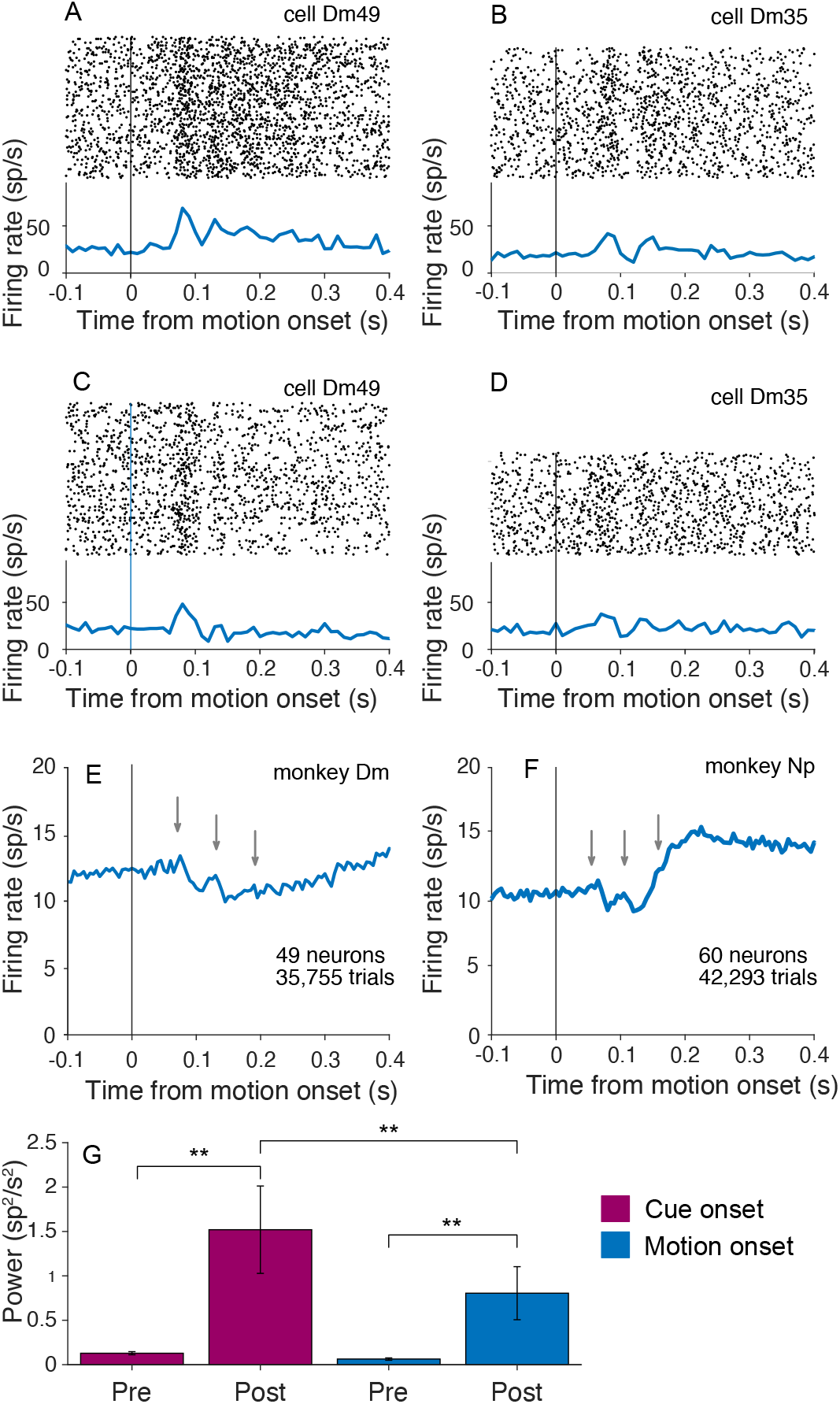
Oscillations at motion onset in the cued attention task. **A–D**, Activity of the neurons shown in Fig. 4A–D, aligned here to the onset of random dot motion. Attention was cued to the upper patch in *A* and *B* and to the lower patch in *C* and *D.* **E,F**, Average activity across all neurons in both locations. Gray arrows mark the positions of the peaks in activity in Fig. 4E&F. **G**, Average 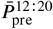 and 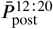, across all neurons from both monkeys, in 90 ms epochs before and after onset of the attention cue and random dot motion stimulus. Asterisks indicate significant differences (*,**: *α* =0.05, 0.01;error bars are s.e.m.).

We also detected oscillations in the local field potential recordings made from the same electrode used for the neural recordings. The LFPs revealed oscillations similar to those detected in the spiking activity. For example, Fig. 6A shows the average LFP for monkey Dm, aligned to the onset of the attention cue. The gray arrows are copies of the black arrows in Fig. 4, which show the peaks in the firing rate oscillations from the same experiments. The oscillations in the LFP recordings from monkey Np are less pronounced, but some deflection is evident at the time of the peaks in spike rate, shown by the gray arrows in Fig. 6B. For both monkeys, 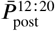 is greater than 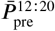 for both cue and motion onset (*p* < 0.0001), and like the firing rate oscillations, 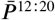 at motion onset is weaker than at cue onset (*p* < 0.0001, Fig. 6C).

**Figure 6.**
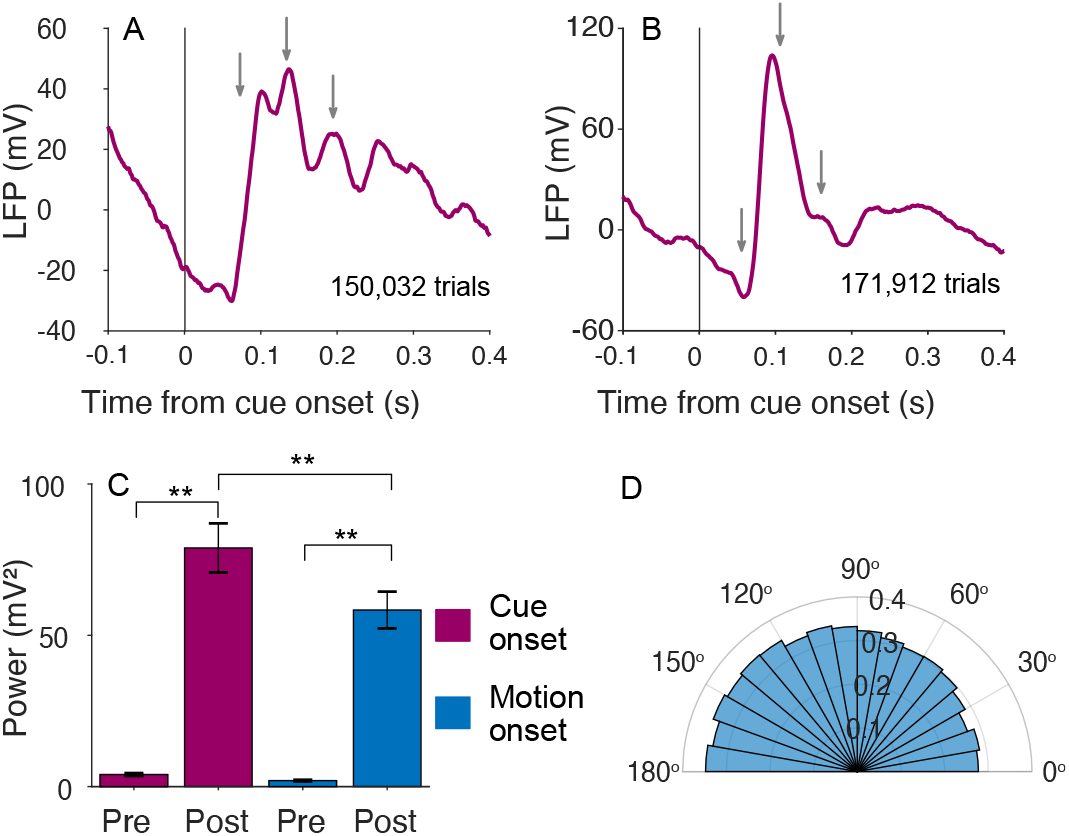
Oscillations are present in the local field potential. **A,B**, Average baseline-corrected LFP from monkeys Dm and Ne, respectively, aligned to onset of the attention cue in the cued attention task. Gray arrows mark the peaks in firing rate activity shown in Fig. 4E & F. **C**, Average 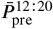 and 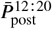, across all sites and both monkeys, measured in 90 ms epochs preceding and following onset of the attention cue and random dot motion. Same conventions as Fig. 5. **D**, Phase alignment of spikes with LFP. Histogram shows the number of spikes in 3°° bins of phase. The phases refer to the cosine carrier of the best Gabor. There is a tendency for spikes to occur more frequently near the trough of the LFP (see Methods, Spike-field alignment).

The oscillations in the LFP and firing rates appear to be manifestations of a common underlying mechanism. In addition to the similarity in their timing and frequency, there is a tendency for spikes to occur in the trough of the LFP oscillation (*p* < 0.0005; Fig. 6D). The frequency histogram of spike phases is obtained by extracting the dominant Gabor atom from the matching pursuit analysis of the LFP. The spike phases are the inverse cosine of the carrier at the time of the spike, such that zero and *π* are the peak and nadir of the carrier, respectively (see Methods, Spike-field alignment). The observation is unsurprising given the similarity of the signals, but it is not an artifact of recording the LFP and action potentials from the same electrode. Similar oscillations in the LFP are present at electrodes that pick up few (or zero) spikes.

### Oscillations in the variable location task

In the variable location task, it is the appearance of the random dot motion itself that resolves the uncertainty about the source of evidence bearing on the decision. As in the cued attention task, the routing must be established between direction selective neurons in the visual cortex that represent the motion and the LIP neurons that represent one of the choice-targets. Here however, connectivity must be established between the onset latency of visual cortical neurons and the beginning of evidence accumulation—roughly40–200 ms from motion onset. The example neuron shown in Fig. 7A–B exhibits oscillations in the firing rates similar to those in the cued attention task. They are also evident in the pooled firing rates across 64 neurons from the three monkeys (Fig. 7C and ***Figure 7—figure Supplement 1***). The average 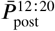 is two orders of magnitude larger than 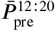 (2.3± 1.2 and 0.034±0.005 sp^2^s^-2^, respectively; *p* < 0.0001).

**Figure 7.**
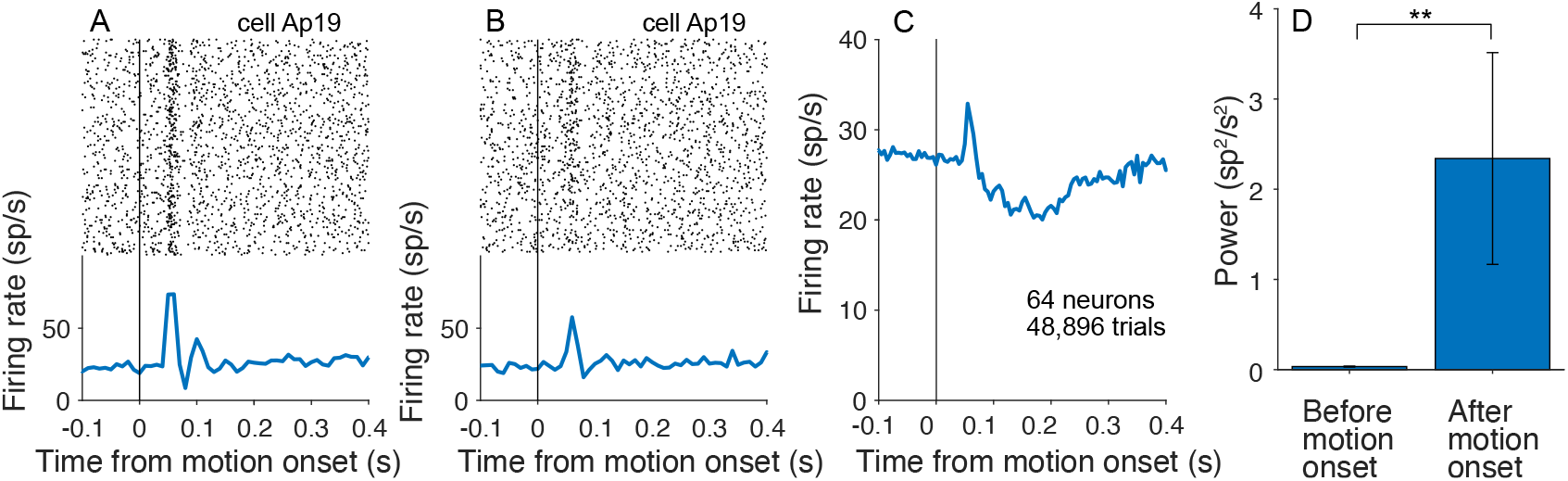
Oscillations in the variable location task. **A**, Activity of an example neuron with stimulus presented above fixation. *Top,* Raster plot of spike times relative to onset of random dot motion. *Bottom* Average firing rates across trials (computed in 10 ms bins). **B**, Same neuron on trials when the motion stimulus appeared below fixation. **C**, Average firing rate, across all neurons from three monkeys. **D**, Average 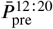 and 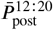, across neurons, measured in 90 ms epochs preceding and following onset of random dot motion. Same conventions as Fig. 5G. **Figure 7—figure supplement 1.** Oscillations in the variable location task for each monkey

The task comprises alternating blocks of fixed and variable stimulus locations. In the former case, it may not be necessary to establish the appropriate functional connectivity on every trial. We therefore predicted that signals associated with routing might be diminished in these blocks. Indeed 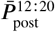 was slightly reduced on trials where the motion stimulus location was fixed (*p* < 0.01, permutation test). While significant, there are aspects of the design that might weaken this comparison. In particular the monkey had learned to expect the RDM to appear in various locations, and may not have adapted fully to the blocked design. We therefore augmented this analysis with a reänalysis of two older data sets, which are better suited to test our hypothesis.

Two monkeys reported in ***Roitman and Shadlen (2002)*** were trained and studied with random dot motion viewed at the center of the visual field. One year later, the same monkeys were retrained and studied on a variable location task (included in ***Shushruth et al. 2018***). We sought to determine whether oscillations in the firing rates were present following onset of the random dot motion. As shown in Fig. 8, we did not detect oscillations in the recordings from either monkey in the earlier *fixed location* study, whereas they are clearly present in the data from same monkeys— and same LIP—in the variable location design (Table 3E). While the study was not designed with this longitudinal comparison in mind, it provides support for the hypothesis that the transient oscillations are associated with neural mechanisms responsible for flexible routing. It also rebuts the assertion that the oscillations are triggered by any task-relevant visual stimulus. The oscillations appear to be associated with task events that resolve uncertainty about the source of information.

**Figure 8.**
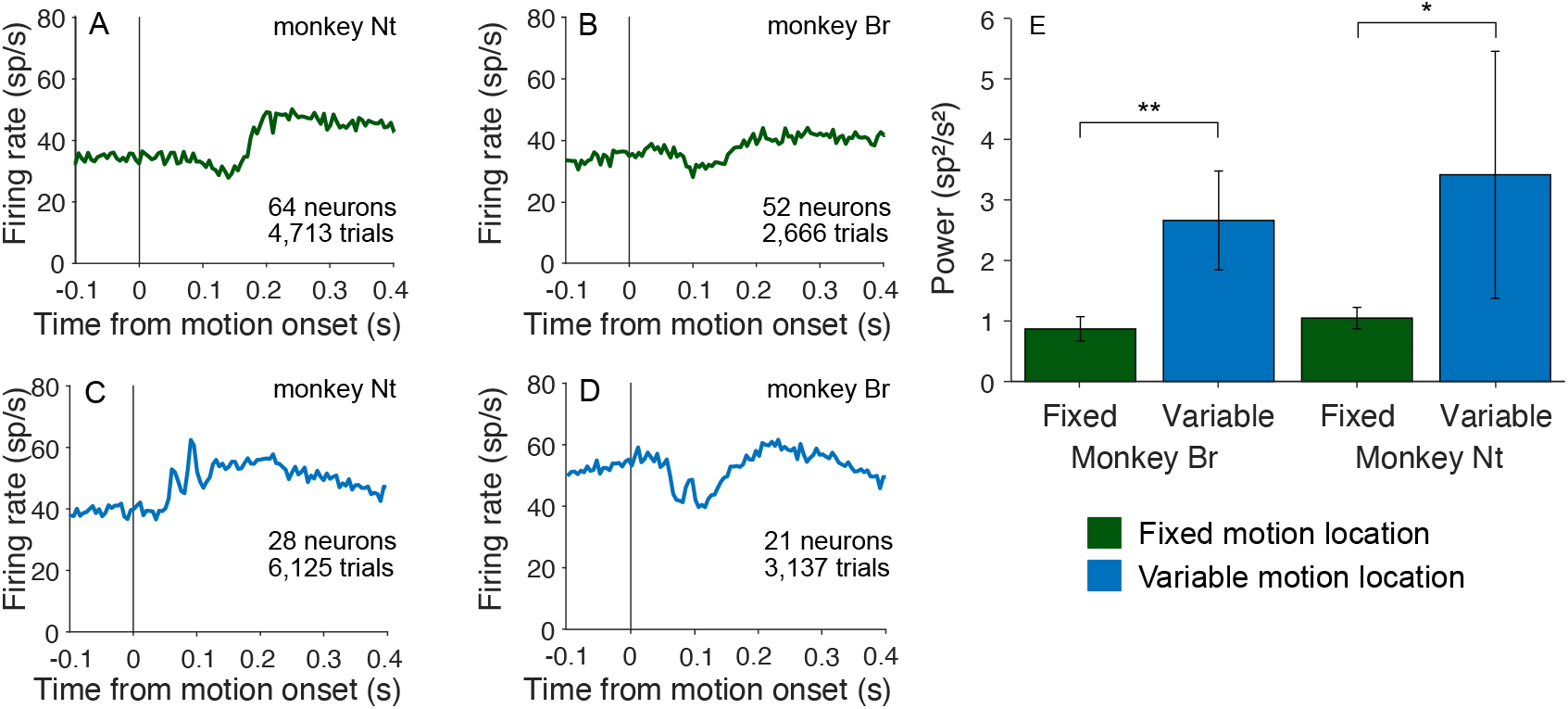
Oscillations emerge after encountering conditions necessitating flexible routing. Two monkeys were first trained and tested with random dot motion stimuli, presented at one central viewing location *(fixed location).* They were subsequently trained and tested with stimuli presented at a variety of locations *(variable location).* **A,B**, Average firing rate aligned to motion onset in the fixed location experiments. **C,D**, Average firing aligned to motion onset in the variable location experiments. **E**, Comparison of Average 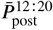 in the two conditions, by monkey. Same conventions as in previous bar graphs.

**Table 3.** Oscillation power in fixed and variable stimulus location tasks. Units are sp^2^s^-2^ (FR, free response task).

|  | Fixed location (FR) |  |  | Variable location (FR) |  |  |
| --- | --- | --- | --- | --- | --- | --- |
| | $\bar{P}_{pre}^{12:20}$ | $\bar{P}_{post}^{12:20}$ | P-value | $\bar{P}_{pre}^{12:20}$ | $\bar{P}_{post}^{12:20}$ | P-value |
| Nt | $0.8 \pm 0.2$ | $1.0 \pm 0.2$ | 0.06 | $0.3 \pm 0.1$ | $3.4 \pm 2.0$ | 0.04 |
| Br | $0.8 \pm 0.1$ | $0.9 \pm 0.2$ | 0.61 | $0.2 \pm 0.1$ | $2.2 \pm 1.0$ | 0.0007 |
| Combined | $0.88 \pm 0.12$ | $0.97 \pm 0.13$ | 0.11 | $0.40 \pm 0.10$ | $2.6 \pm 0.9$ | 0.00001 |

Of course, routing requires specification of both the source and destination of information. We therefore looked for oscillations following the onset of the choice targets. Recall that this event precedes the attention cue in the cued attention task and the the RDM in the variable location task. As shown in Fig. 9, oscillations are present in both tasks following onset of the choice targets (*p* < 0.0001), one of which is in the neural response field. They can also arise when both targets appear outside the neural response field, but are relevant to the routing of other information in the response field. This occurs in the second of the reänalyzed data sets, in blocks where the RDM is displayed in the neural response field (0.21 ± 0.07 and 0.54± 0.11 sp^2^s^-2^ for 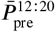 and 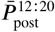, respectively; *p* < 0.001). This observation, like the attention cue in Fig. 4, is another example of an oscillation caused by the onset of stimuli outside the neural response field, but relevant to neurons that represent the predicted retinotopic location of the motion evidence.

**Figure 9.**
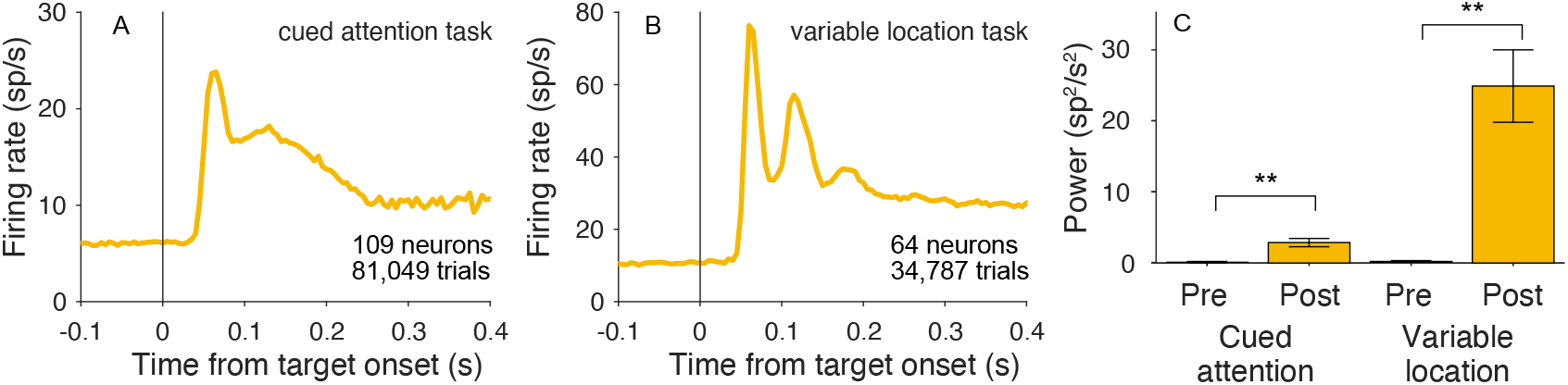
Oscillations are present at target onset. **A**, Average firing rate aligned to onset of the choice targets for both monkeys in the cued attention task. **B** Average firing rates aligned to onset of the choice targets for all three monkeys in the variable location task. **C**, Average 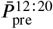 and 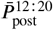 across neurons, measured in 90 ms epochs preceding and following onset of targets. Same conventions as in previous bar graphs.

## Discussion

Unlike innate sensory-response programs, such as escape or courtship, evolution did not imbue the brain with circuits devoted to the vast repertoire of decisions one encounters in life—including the motion tasks studied here. Whereas the processing of motion and the organization of orienting eye movements rely on dedicated sensory and motor circuits, the neural circuits responsible for planning possible eye movements cannot exploit dedicated connections to the neurons that represent all the possible sources of evidence bearing on such plans. The flexibility to learn which sources are relevant and to route them in the moment are hallmarks of higher brain function.

We studied an example of flexible routing by introducing uncertainty about the source of visual evidence bearing on a decision about motion direction. In one task, the location of a single patch of random dots was varied randomly across trials. In the other, an attention cue indicated which of two motion patches should inform the decision. In both tasks the decision is communicated by a saccadic eye movement to one of two choice targets. An advantage of this highly studied perceptual decision is the accompanying quantitative framework that unites choice accuracy, sensitivity, decision time, change of mind, and confidence *(**Shadlen and Kiani, 2013***). We exploited this framework to show that a portion of the errors on the cued attention task are explained by some form of misrouting. Both monkeys fail to suppress all information from the uncued patch, and monkey Dm appears to attend to the wrong patch altogether on 3–6% of trials, accounting for at least half of the errors at the strongest motion coherence. The observations are consistent with a large body of work in cognitive science that frames attention in terms of the control of information flow ***(Driver, 2001; Posner, 1988; Posner et al., 2004; Buschman and Miller, 2009; Buschman and Kastner, 2015; Panichello and Buschman, 2021***). In the present study, this control problem, what we refer to as routing, must be completed by the time neurons in association cortex begin to integrate the sensory evidence toward a decision. The destination of the routed information is specified when the choice targets appear, but the source is uncertain until the onset of motion in the variable location task and the onset of the attention cue in the cued attention task.

We discovered a neural correlate of this routing event in the lateral intraparietal area (LIP). We focused our recordings on neurons with two properties: *(i)* a response field that overlaps one of the choice-targets and (*ii*) spatially selective persistent firing rates during an oculomotor delayed response task. Such neurons are known to represent the accumulation of the noisy evidence used by the monkey to inform the saccadic choice, and we replicated this phenomenon (Fig. 3). We observed that many such neurons also exhibit a brief oscillation in firing rate that is time-locked to the moment when information about the source of evidence becomes available. The oscillation manifests as a transient excess of spikes that repeats one or more times at intervals of 61 ± 2 ms (~16 Hz), and it appears to be coupled to the local field potential (Fig. 6). Striking examples like those in Figs. 4, 5 and 7 were rare, but the majority of cells showed an increase in oscillatory activity in this range, but they were undetectable in the data from ***Roitman and Shadlen (2002)***, who presented the motion stimulus in the same location on all trials. The oscillations appeared in the same monkeys (and same recording sites) after they were trained to base their decisions on stimuli that could appear in different locations (Fig. 8). This serendipitous observation supports the hypothesis that the phenomenon is associated with flexible routing of information from neurons that represent the stimulus motion to neurons in LIP that represent the decision. The connection path is almost certainly polysynaptic.

Previous studies have identified transient oscillations, also in the low beta range, that corre-late with performance on perceptual tasks (***Koelewijn et al., 2008**; **Haegens et al., 2011**; **Siegel et al., 2011***). For example, ***Donner et al. (2007)*** describe such oscillations originating from posterior parietal cortex following onset of a random dot motion stimulus. They reported that power was greater when a stimulus was correctly categorized as motion or noise (hits and correct rejects) than on misses and false alarms. We interrogated all of our data sets for such a relationship but found only one case: a small reduction in 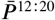 aligned to motion onset on errors in the cued attention task (*p* < 0.006, permutation test). Other than this one case, we did not detect a convincing correlation between behavioral performance on the motion task and the magnitude of the oscillations. Perhaps we lack sufficient power. However it seems more likely that errors were not associated with a failure to route, but rather, a failure to route the right information. Consistent with this interpretation, ***Fiebelkorn et al. (2013)*** reported periodicity in detection accuracy during visual detection tasks. The periodicity was synchronized with cycles of the theta rhythm measured in the LFP recorded from area LIP, and the poor-detection phases were associated with *increased* power in an associated 10–18 Hz frequency band (***Fiebelkorn et al., 2018, 2019***), which the authors interpret as a sign of attentional shifts away from the task. These shifts in attention might involve the same routing processes as allocation of attention in our task.

Why would oscillations be associated with routing? One possibility is that they serve to syn-chronize spikes and thus increase their influence on downstream circuits *(**Singer and Gray, 1995**; **Akam and Kullmann, 2010; König et al., 1995**; **Gregoriou et al., 2009**; **Fries, 2005,2015; Buschman and Miller, 2009***). If so, they ought to be present during the epoch in which signals in upstream motion areas are affecting the LIP response. In our data, however, they are present only tran-siently, in the epoch preceding the transfer of information (cf. ***Panichello and Buschman 2021***). We therefore infer that they are associated with the mechanism that establishes the connection rather than facilitating the flow of information directly. Of course, the oscillations themselves do not form the connections, but they may provide a clue to the underlying mechanism. Among the many challenges posed by routing is the need to identify the appropriate neurons at the source and destination. We suspect that the oscillatory signal is linked to this identification function.

It has been shown that field potentials (e.g., eCoG) are associated with calcium *plateau* potentials in apical dendrites of layer-5 pyramidal neurons (***Suzuki and Larkum, 2017***), and these same potentials are capable of inducing plastic changes at relevant time scales (e.g., behavioral time scale plasticity, ***Magee and Grienberger 2020***). Such plateau potentials and their biochemical sequelae might allow long range projections—especially feedback—to identify their targets, or for the targets of the projections to establish a state of receptivity to a signal that is broadcast widely (***Quinn et al., 2021***). This would be a convenient way for feedback projections to pick out the causes of the activity that is feeding back. This might serve many functions, including learning to use those inputs again under the right conditions, or to bind in some way the cause of an event with with its consequences. Oscillatory activity might be a signature of these inputs (***Zhang and Bruno, 2019***). Thus we are in agreement with a longstanding view that oscillations, measured mainly in the field potentials, herald a connection to cognitive states that firing-rates alone do not divulge (***Freeman et al., 1983; Singer and Gray, 1995; Fries et al., 2001; Crick and Koch, 2003***).

## Methods and materials

Four adult male rhesus monkeys *(Macaca mulatta) were* implanted with a titanium headpost (Rogue Research, Montreal, Canada) and a plastic (Peek) recording chamber (Crist Instruments, Damascus, MD). The placement of the chamber was guided by 3D reconstruction of MRI scans (OsiriX DICOM Viewer, Pixmeo, Bernex, Switzerland) to ensure access to area LIP along the left intraparietal sulcus. In the experiments, the monkeys were seated in a primate chair (Crist Instruments) that was custom fit to support the monkey’s size and weight during head stabilization, allowing the monkey to adjust its posture below the head and thus prevent potential discomfort associated with head stabilization. Extracellular single-neuron recordings were made using quartz coated tungsten microelectrodes (Thomas Recording GmbH, Giessen, Germany) or 16-channel V-probes or S-probes (Plexon), which were advanced (Mini Matrix drive, Thomas Recording) through a metal guide tube seated in a plastic grid. Electrical recordings were filtered and amplified (Ominplex recording system, Plexon Inc, Dallas, TX). Waveforms identified as single neuron action potentials were saved, and each occurrence was assigned a spike-time. The quality of isolation was confirmed offline based on interspike interval and clustering based on principal component analysis of the waveforms (Plexon Offline Sorter, Plexon Inc, Dallas, TX).

All procedures were approved by the Columbia University IACUC and conform to the NIH guide for the care and use of laboratory animals (***National Research Council, 2011***).

### Behavioral tasks

Monkeys were trained to perform a variety of oculomotor and perceptual tasks that required the monkey to maintain the gaze on a fixation point and to make saccadic eye movements to visual targets in the periphery (see Table 2). Eye position (gaze angle) was measured with high speed video tracking (EyeLink 1000, SR research). Acceptance windows for eye position during fixation was a square ± 1.5° from the fixation point (i.e., 9 deg^2^). Here and throughout, °, or *deg,* stands for degrees visual angle. For saccades to peripheral targets, the acceptance window was a ± 5° square around the target center. The criteria were relaxed for eccentricities exceeding 12°.

#### Cued attention task

Two monkeys were trained to perform a variation of a random dot motion direction-discrimination task used in previous studies (e.g., see ***Roitman and Shadlen 2002***) in which two motion patches were shown, but only one was informative. In this cued attention task (Fig. 1) the monkey initiated a trial by fixating on a central red dot. After 0.35 s two white targets appeared, each with a diameter of 0.5 degrees visual angle. Target onset was followed by a delay period, drawn from a truncated exponential distribution

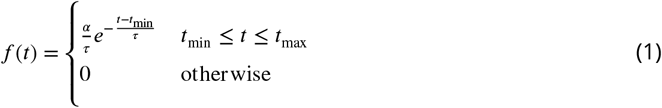

where *τ* = 0.1, *t*_min_ = 0.2 s, *t*_max_ = 0.6 s, and *α* is chosen to ensure the total probability is unity. Note that the expectation of *t* is less than *t*_min_ + *τ*, owing to truncation. In what follows all variable delay periods are described by a range, *t*_min_ to *t*_max_, and *τ* in Eq. 1.

The cue, a 5° blue ring, was flashed on the screen 4.5° directly above or below the fixation point for 0.18–1 s (*τ* = 0.2 s). After a 0–1 s delay (*τ* = 0.15 s), two motion patches appeared. The monkey was required to attend to the cued motion patch while ignoring the irrelevant motion in the uncued stimulus. The direction of motion of the uncued stimulus was the same as that of the cued stimulus on one third of trials and opposite on two thirds. We found this ratio worked to minimize the influence of the uncued motion stimulus on the choices. Motion was always along the horizontal axis. For each trial, the strength of the net motion (motion coherence) was drawn uniformly from the set{0,3.2,6.4,12.8,25.6, 51.2}. The strength of motion for the two patches was matched to prevent the monkey from responding based on the easier motion stimulus ratherthan the relevant one. The cued location was assigned randomly between the two locations for each trial. Motion lasted for 0.1–1.5 s (*τ* = 0.35 s) and was followed by a 0.4–1.2 s delay (*τ* = 0.15 s). After this delay the red fixation point disappeared and the monkey indicated its response by a saccade to a left or right choice target.

One choice target was centered in the response field of the recorded neuron. The other was at the same elevation and azimuth in the opposite hemifield. The random dot motion was confined to two circular apertures (5° diameter) centered at the same elevation above and/or below the fixation point. The locations were determined by establishing the extent of the neural response field so as to avoid overlap. The monkey performed a series of delayed saccades (see Mapping tasks), and we ensured that saccadic targets (white spots, 0.5° diameter) did not elicit a visual or memory response when they overlapped the intended apertures.

#### Variable location task

In the variable location task, only one of the random dot patches was shown on the trial, and there was no attention cue. The same strategy was employed to determine the two locations, above and below the fixation point, but the the location on any trial was random (Bernoulli dist, P=0.5). For monkeys Ap and Dz, the choice targets were not restricted to the same elevation in the visual field. One was centered in the neural response field (location *A*); and the other was at location *B,* such that a virtual line *AB* passes through the fixation point (*F*), and *AF* ≅ *BF*). The opposing directions of motion were parallel to *AB*. Monkey Dm was only trained on horizontal motion, so targets and motion shared the same elevation. The task was otherwise similar to the cued attention task.

For monkeys Ap and Dm, motion was displayed for 0.1–1 s (Eq. 1, *τ* = 0.25 s) and was followed by a 0.4–1.2 s delay (*τ* = 0.15 s). After this delay the fixation point disappeared and the monkey indicated its decision by making a saccade to one of the choice targets. Motion strengths were drawn from the same distribution as in the cued attention task for monkeys Dm and Ap; the strongest coherence was not included for monkey Dz. For this monkey we used a free response (choice-response time) design. the task was identical to the controlled duration version, except that a saccadic response was accepted any time after motion onset. For all monkeys, correct choices were rewarded with a drop of juice. Trials with 0% coh were rewarded randomly. Experiments were conducted in alternating blocks of 120 trials with either a fixed or variable stimulus location. In a fixed location block the motion stimulus appeared in one location for 60 consecutive trials and then appeared in the other location for 60 trials.

For monkeys Ap and Dz stimuli were shown on a 40 cm cathode-ray tube (CRT) monitor with a 75frame/second refresh rate. For monkey Dm, stimuli were shown on a 54 cm liquid crystal display (LCD) with an effective refresh rate of 60 frames per second. For this display, the interval between replotted frames was reduced from every third frame to every second frame. we adjusted the dot displacement to achieve consistent speed across display types (typically 5°/s).

#### Mapping tasks

We conducted two screening tasks to select neurons for study in the tasks. In both, the monkey maintained its gaze on a central fixation point (FP), and initiated a saccade when the FP was ex-tinguished. In the *memory saccade task,* 0.2–1 s (*τ* = 0.2 s) after attaining central fixation, a white target (0.5° diameter) was flashed in the periphery. After a memory delay of 0.7–1.2 s (*τ* = 0.1.5 s) from target onset, the fixation point was extinguished. disappeared and the monkey was free to saccade to the cued location to receive a juice reward. The *overlap saccade task* was the same, except that the target remained visible throughout the delay period and the saccade. We refer to both these tasks as ‘oculomotor delayed response’ (ODR).

### Neuron selection and recording

Recording sites were selected by 3D reconstruction of anatomical MRI (3T). The electrode was advanced along the intraprietal sulcus at positions that are thought to correspond to the ventral portion of the lateral intraprietal area (LIPv; ***Lewis and Van Essen 2000***) where one encounters meany neurons with visual and perisaccadic responses. Within putatative LIPv, we mapped all well isolated units using the overlap saccade task. Neurons with spatially selective persistent activity in this task were further mapped using the memory-saccade task. A neuron was included in the data set if it showed spatially selective persistent activity during the delay period of memory saccades and if the neural response field allowed for a task geometry compatible with the monkey’s training. We excluded neurons *post hoc* if we obtained less than 240 trials before the signal to noise deteriorated to the point that the spike waveform was not adequately isolated (7/71 neurons).

In the cued attention task, a 16 channel probe was used to record several neurons simultaneously. All channels were screened with the memory guided saccade task. The recording probe was positioned to maximize the number of recorded units showing memory activity. The task objects could not be placed optimally for all cells, but nearby cells tended to have similar response fields. The task geometry was optimized for the best isolated channel. This yielded 1–7 simultaneously recorded cells with acceptable task geometry: a choice target roughly centered in the response field and both motion patches outside the response field. Cells were sorted offline as for single electrodes. Particular attention was paid to whether waveform principal components, or spike rate changed over time to ensure that the same cell was recorded throughout the session. Monkeys performed an ODR trial to each target location after every 40 trials on the motion task, and thorough screening was repeated at the end of the session to ensure that response fields were constant throughout the session. If a cell showed a change in any of these parameters, trials after that change were excluded from analysis. Occasionally a new waveform appeared during the recording session. It was included in the analysis if (*i*) it was well isolated from background noise, (*ii*) exhibited a consistent waveform-principal components, spike rate, and response preference in the interleaved ODR trials, and (*iii*) showed an appropriate response field in the post session screening tasks.

### Data analysis

Peristimulus time histograms (PSTHs) were generated by aligning spike times to an event of interest and finding the average number of spikes, across trials, in time-bins relative to the event. Time-bins were 5 ms wide for averages across neurons and 10 ms for single neurons. For the firing rate vs. time graphs (by coherence) traces in Fig. 3, the rates are obtained by convolving the point process, *δ*(*t* - *s_i_*), where *S_i_* are spike times, with a non-causal boxcar filter of width 100 ms. This smoothing was not applied to any other plot or analysis, as it obscures the oscillations of interest. To better visualize the decision-related activity, we detrended the responses in Fig. 3. For each neuron we subtracted the average response to the 0 and +/-3.2% coherences. Figures show the average across neurons, with each neuron weighted by the number of recorded trials. Across the two tasks, 8 out of 173 neurons showed a preference for the ipsilateral direction during the motion viewing epoch. For these neurons, the sign of the motion was reversed in analysis of signed coherence (Fig. 3).

#### Behavior

The decision process leading to leftward and rightward choices is affected by the direction and strength of motion as well as the duration of the stimulus. The durations were controlled by the experimenter for the three monkeys displayed in (Fig. 2). For the fourth monkey, Dz, we used a free response (choice-response time) design. Decision formation in both designs is explained by a process of bounded accumulation of noisy evidence, also known as bounded drift-diffusion ***(Kiani et al., 2008***). Accordingly, momentary motion evidence is integrated over time until it reaches one of two bounds (±*B*) or the evidence stream is turned off. The influence of the motion evidence depends on the signed motion coherence (*C*) and on a drift rate parameter (*κ*).

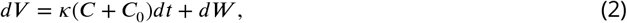

where ***W*** is a standard Wiener process (i.e., ***dW*** is a sample drawn from a Normal distribution, 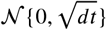). The initial state is ***V***_*t*=0_ = 0 and the process continues until |***V***(*t*)| ≥ ***B***. The time of this termination governs the response time in a free response task (e.g., monkey Dz), and simply curtails further integration when the stimulus duration is controlled experimentally. If the decision process is terminated when integrated evidence reaches a bound, the chosen direction is the sign of the bound reached. If a bound has not yet been reached before the evidence stream is turned off (at *t* = *t*_dur_) the chosen direction depends on the sign of the unabsorbed integrated evidence. The choice probability was modeled by fitting ***B**, κ*, and a bias term *C*_0_ expressed as an offset in signed motion coherence *(**Hanks et al., 2011; Urai et al., 2019***). These quantities are obtained by numerical solution of the Fokker-Planck equation, Which yields a probability density comprising three components: (*i*) *f*_+_(*t*\*t* ≤ *f*_dur_), the upper bound absorption times, (*ii*) *f*___ [*t*|*t* ≤ *t*_dur_), the lower bound absorption times and (*iii*) *f*_un_(***V**/t* = *t*_dur_) the values of the unabsorbed ***V*** at *t* = *t*_dur_, such that

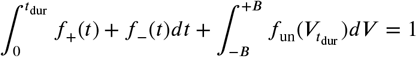

The probability of a positive choice is

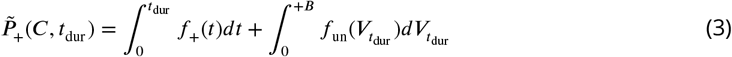

and

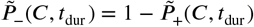

This specifies the *base* diffusion model without misrouting. The latter comprises (1) attention to the wrong patch and (2) incomplete suppression of the uncued motion patch. For the base model,the observed proportion of positive choices is 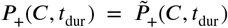. If the monkey attends to the wrong motion patch on a fraction of trials, *λ,* then

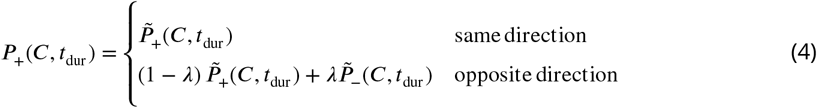

To model incomplete suppression of the uncued patch, we allow for a different value of *κ* in Eq. 2 when the patches have the same or opposite directions

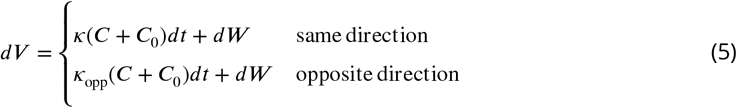

where *C* is the signed coherence of the cued patch.

The models were fit separately for the two monkeys using maximum likelihood. The fitted parameters are {*κ,C*_0_,***B***} for the basic model without misrouting (df=3). Erroneous routing and incomplete suppression add one degree of freedom apiece. We report the absolute value of the ΔBIC to convey support of a model against an alternative (i.e, Bayes Factor >1; Table 1).

#### Quantification of oscillations

We implemented a *matching pursuit* (MP) algorithm to quantify the strength of oscillations in the neural firing rates and local field potentials *(**Chandran et al., 2016; Mallat and Zhang, 1993***). MP is a greedy algorithm designed to represent a finite signal, *s*(*t*), as a sum of Gabor functions (atoms) from a library that covers the position *τ* and width *σ* of the Gaussian envelope as well as the angular frequency *ξ* of the carrier sinusoids:

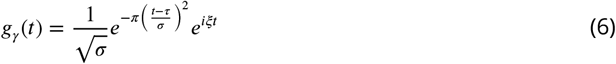

where the subscript, *γ*, identifies the atom, *γ* = {*τ,σ*,}. MP is well suited to brief epochs containing mixtures of transient and periodic features. We used the open source algorithm developed by the Epilepsy Research Laboratory at Johns Hopkins Medical Institutions and Supratim Ray (available from https://github.com/supratimray/MP). Forthe spike rates, *s*(*t*) is the average firing rates across trials for a neuron, as shown in the example neurons (e.g., using −0.3 ≤ *t* < 0.724 s relative to the event of interest. For spiking data, the input is the averaged unsmoothed firing rate (1 ms bins). For LFP data, the input is the trial averaged voltage in (1 KHz sampling rate). The output is power as a function of time and frequency, as shown in ***Figure 4—figure Supplement 1***. We define the low-beta power as the mean Wigner-Ville power 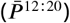 in the frequency band 12–20 Hz in the 90 ms before or 40–130 ms after event onset, denoted 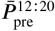 and 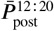, respectively. We typically report the mean 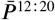 across neurons (± s.e.m.) and determine statistical significance by applying a Wilcoxon signed-rank test (a nonparametric equivalent of the paired t-test), using 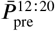 and 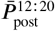 for each neuron. For comparisons of unpaired 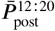, we use the Mann-Whitney *U* test.

The estimate of 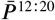 can be impacted by the number of samples in the mean firing rate or LFP. For comparisons between conditions with unequal numbers of trials, we also evaluated the mean difference in 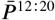 derived from random subsets of *W*æ trials from the two conditions, where *N*_10_ is ~ 10% of the number of trials in the condition with the lesser number of trials. We calculated 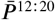 from the spike rate averages in the two conditions and took the difference, *D*_pow_. We repeated this procedure 1000 times and used the mean, 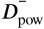 as the estimate. We compared this test statistic to its distribution under the null hypothesis, by repeating the identical procedure on random subsets drawn from the union of the data from the two conditions, again using 1000 repetitions to achieve a sample of 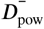 under the null hypothesis. We repeated this 100 times to estimate its distribution, and calculated the p-values from the tail probabilities (2-tailed). This bootstrap procedure produces qualitatively similar results to those obtained from the Mann-Whitney *U* in almost all cases (e.g., comparison of 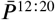 triggered by motion onset in the old data sets). The two exceptions are the comparison of 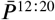 after motion onset on correct vs. errors and on blocks of variable location and fixed location of the RDM. This is why we qualify our interpretation of these findings.

For single neuron analyses, the data from each neuron was divided into 50 trial blocks. For each block we obtain 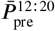 and 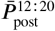 and applied a Wilcoxon signed rank test to evaluate the null hypothesis of identical means.

#### Cell type analysis

Spike waveforms were preserved for 42 neurons from monkey Np. Neurons were classified as putative excitatory or inhibitory based on the distance, *D* between the peak and trough of the average spike waveforms. All but two were classified as either putative inhibitory *(D ≤* 150 μs) or excitatory (*D* ≥ 350 μs; ***Barthó et al. 2004***; **Trainito et al. 2019**; **Ardid et al. 2015**).

#### Spike-field alignment

To assess the relationship between the oscillations in firing rate and local field potential, we estimated the phase of the LFP associated with all spikes that occur in an epoch 40–130 ms after cue onset. The analysis was restricted to neuron-LFP recordings where the MP algorithm identified oscillations in the LFP (31 of 104 experiments; cued attention task), based on the criterion that at least one of the 10 strongest atoms (Gabor functions) overlapped the epoch and frequency band of interest (i.e., 12–20 Hz), based on its carrier. We used the inverse cosine of the latter to associate the time of each spike with a phase. Thus spikes occurring near the peak or trough of the oscillation are assigned phases *ϕ_s_* ≈ 0 and *ϕ_s_* ≈ *π*, respectively. To produce the histogram of phase values in Fig. 6D, we correct for the non-uniform representation of cosine phase in the sampled epochs. We evaluated the null hypothesis that *ϕ_s_* is uniformly distributed by comparing the empirical (non-uniform) representation of candidate phases with the distribution of *ϕ_s_* (Kolmogorov-Smirnov two sample test).

## Acknowledgments

We thank Chris Fetsch for piloting initial experiments; Cornel Duhaney and Brian Madeira for exceptional assistance in animal training and care; Supratim Ray for advice on implementation of the Matching Pursuit algorithm; Danique Jeurissen, Shushruth, Natalie Steinemann, and Gabe Stine for comments on an earlier draft of the manuscript.

---

**Figure 3—figure supplement 1.**
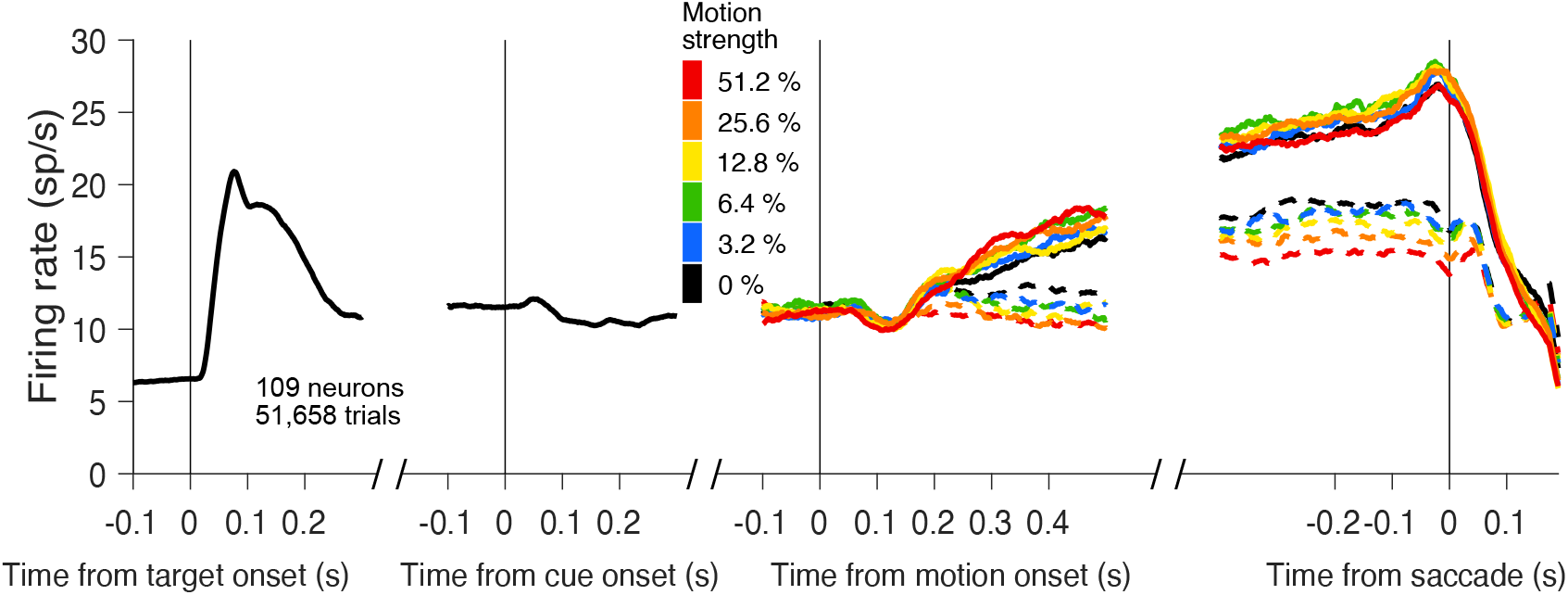
Firing rates aligned to all task relevant events. These are the same data in Fig. 3. From left to right, responses are aligned to target onset, cue onset, motion onset, and saccade. For motion and saccade epochs, warmer colors indicate stronger motion. Solid and dashed traces indicate choices to the target in and out of the response field, respectively.

**Figure 3—figure supplement 2.**
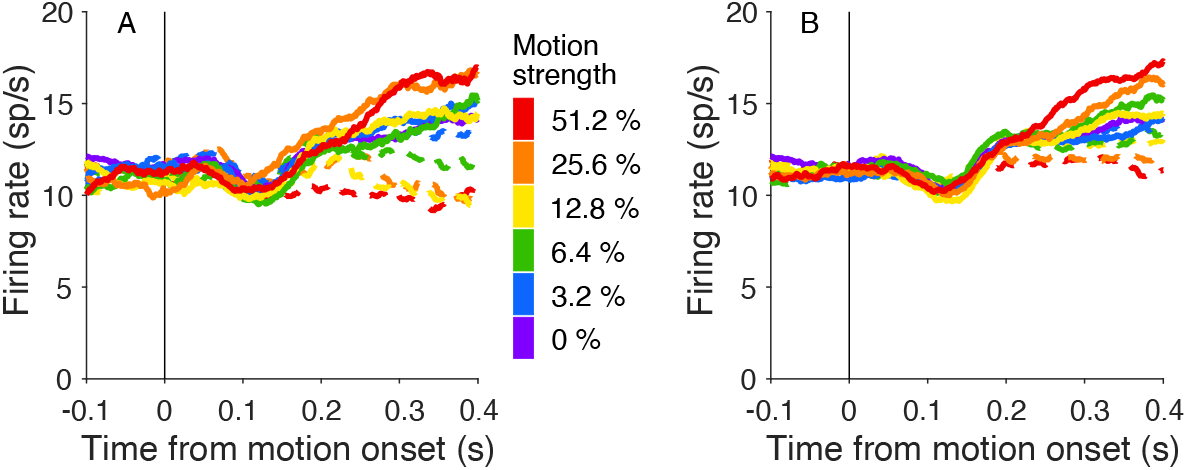
Comparison of responses when motion patches had the same or opposite directions. The graphs use the combined data from both monkeys without detrending, including error trials. **A**, LIP activity for trials with motion in the same direction in both patches. Firing rates are averages using all trials with the same motion strength (color) and direction (line style). The responses begin to exhibit a dependency on motion direction and strength ~ 180 ms after motion onset. **B**, LIP activity for trials with opposite directions of motion in the two patches. Activity follows the same general pattern seen in *A,* but the dependency on motion strength is less apparent.

**Figure 3—figure supplement 3.**
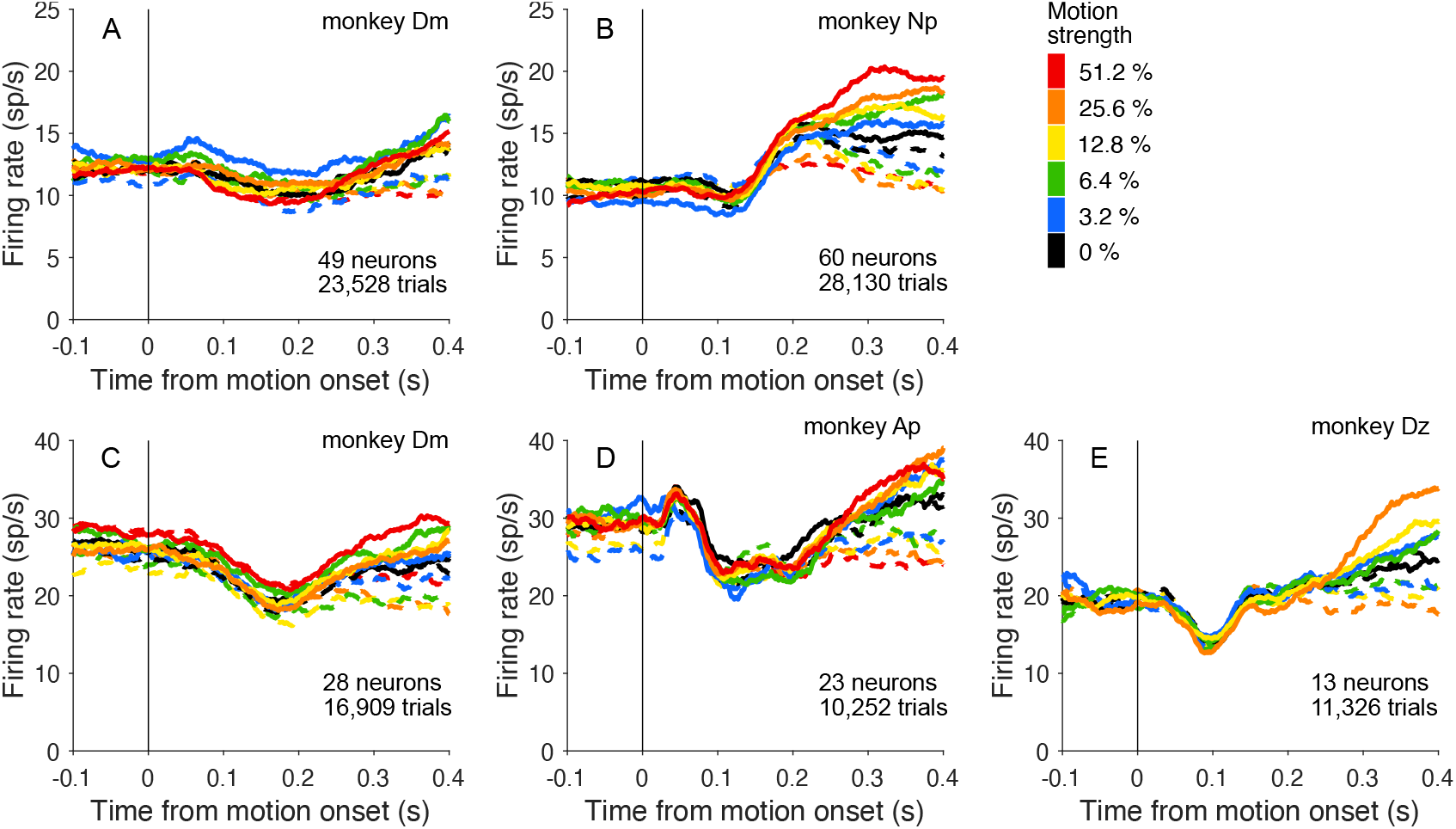
Neural response shown separately for each monkey and task. Data are from the same neurons included in Fig. 3 shown here without detrending. **A,B**, Average firing rate aligned to motion onset in the cued attention task. **C–E**, Average firing rate aligned to motion onset in the variable location task.

**Figure 4—figure supplement 1.**
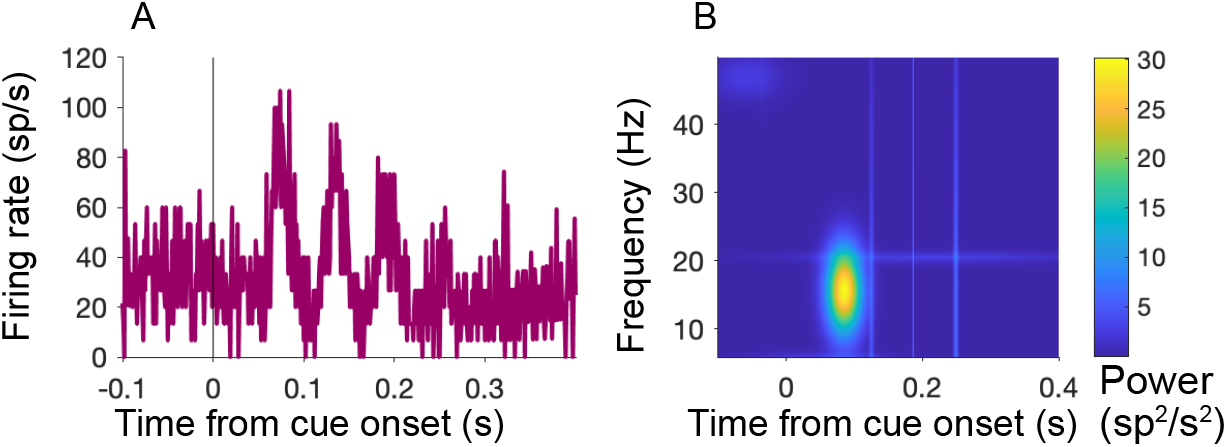
Oscillations in spiking activity measured using a matching pursuit algorithm. The algorithm uses a greedy method to fit the waveform using a dictionary of Gabor functions of time (Eq. 6). **A**, Input to matching pursuit algorithm. The average firing rate is rendered as a peristimulus aligned histogram (1 ms bin width) from neuron Dm49 aligned to cue onset (same 150 shown in Fig. 4A). **B**, Output of matching pursuit algorithm. Heat map shows power (color) by frequency and time from cue onset.

**Figure 4—figure supplement 2.**
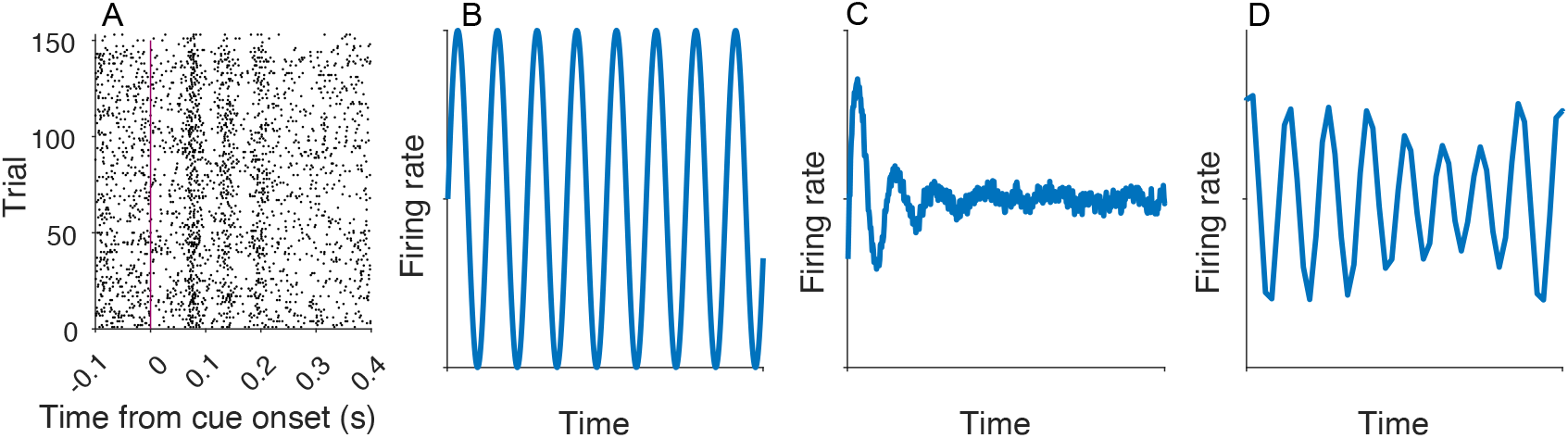
Realigning does not identify additional peaks. **A**, Activity realigned using the affinewarp algorithm (***Williams et al., 2020***). **B–D**, Affinewarp can recover temporally jittered oscillations in synthetic data. **B**, Input oscillation. **C**, Mean activity of 500 simulated trials using the firing from *B,* with increasing temporal jitter. For each trial temporal noise is added at each time point before generating spikes, causing the oscillations to become misaligned in time and to disappear from the average. **D**, Average firing rate of the activity of the trials shown in *C* after applying the affinewarp algorithm. The underlying oscillation is partially recovered.

**Figure 7—figure supplement 1.**
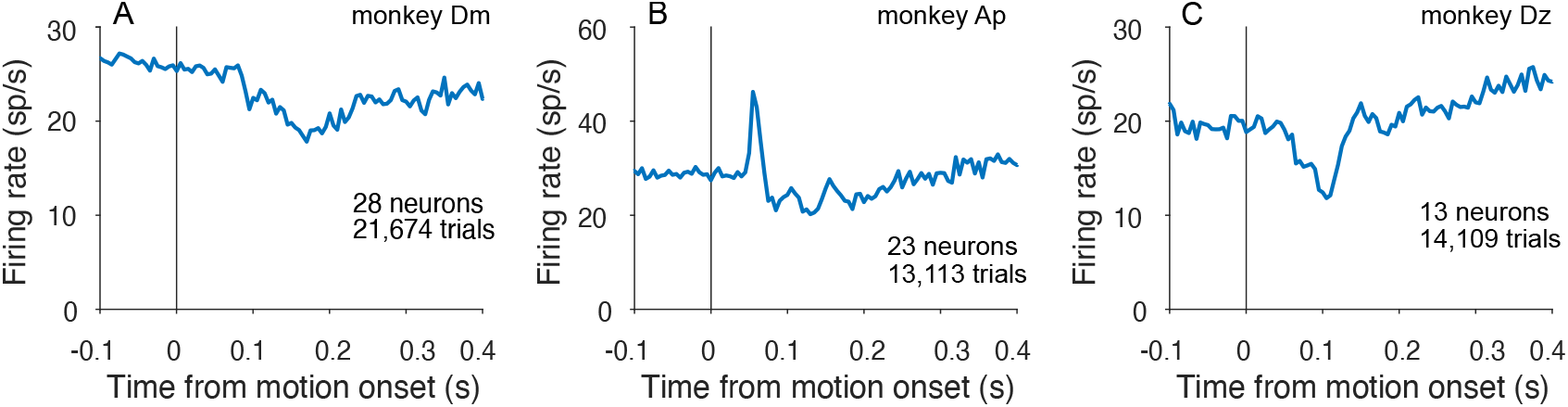
Oscillations in the variable location task for each monkey. Average firing rates, aligned to motion onset.

## Notes

### Competing Interest Statement

The authors have declared no competing interest.

### Summary of Updates

Minor changes to Introduction; clarifications on the use of the Matching Pursuit algorithm.

